# A wrappable microwire electrode for awake, chronic interfacing with small diameter autonomic peripheral nerves

**DOI:** 10.1101/402925

**Authors:** Jessica D. Falcone, Tristan Liu, Laura Goldman, David D. Pogue, Malgorzata Straka, Loren Rieth, Chad E. Bouton, Harbaljit S. Sohal

## Abstract

Bioelectronic medicine requires the ability to monitor and modulate nerve activity in awake patients over time. The vagus nerve is a promising stimulation target, and preclinical models often use mice. However, an awake, chronic mouse vagus nerve interface has yet to be demonstrated. Here, we developed a functional wrappable microwire electrode to chronically interface with the small diameter mouse cervical vagus nerve (∼100 μm). In an acute setting, the wrappable microwire had similar recording performance to commercially available electrodes. A chronic, awake mouse model was then developed to record spontaneous compound action potentials (CAPs). Viable signal-to-noise ratios (SNRs) were obtained from the wrappable microwires between 30 and 60 days (n = 8). Weekly impedance measurements showed no correlation between SNR or time. The wrappable microwires successfully interfaced with small diameter nerves and has been validated in a chronic, awake preclinical model, which can better facilitate clinical translation for bioelectronic medicine.

---

With recent advancements in bioelectronic medicine, electrical stimulation of the vagus nerve has become a clinical therapy for epilepsy(Morris & Mueller 1999; Morris et al. 2013; Handforth et al. 1998; George et al. 1995) and depression (Nahas et al. 2005; Sackeim et al. 2001; Rush et al. 2005), and in recent years, has shown efficacy in treating inflammatory auto-immune diseases (D’Haens et al. 2018; Bonaz et al. 2016; Koopman et al. 2016). Preclinical murine research has demonstrated the impact of acute vagus nerve stimulation on modulating the immune system (Inoue et al. 2016; Borovikova et al. 2000; de Jonge et al. 2005; Rosas-Ballina et al. 2011; Patel et al. 2017). By further studying the neural activity of the vagus nerve, real time recordings have the potential to be used as a therapeutic marker to monitor inflammation (Silverman et al. 2018; Steinberg et al. 2016) and to evaluate stimulation efficacy (Bouton et al. 2016; Bouton 2017; Bouton & Czura 2018). To better understand disease pathways and stimulation paradigms, the development of a chronic, awake rodent model for recording from the vagus nerve is critical.

The primary interfacing technology for the vagus nerve are cuff-based electrodes, which have also received FDA approval and are marketed for stimulation to treat epilepsy and depression (TERRY et al. 1991; Krahl 2012). Other clinical electrodes already exist for stimulating and recording from peripheral nerves (Micera et al. 2011; Kundu et al. 2014; Ledbetter et al. 2013; Schiefer et al. 2010); however, these electrodes penetrate and manipulate the nerve, and the safety and efficacy of these approaches needs additional study given the autonomic innervations (heart, lungs, etc.) of the vagus nerve. Preclinical studies for chronic vagus nerve electrodes have been successfully conducted in large animal models (Zhang et al. 2009; Ardell et al. 2017; Hamann et al. 2013; Metcalfe et al. 2018; Valdés-Cruz et al. 2002) and in rats (Somann et al. 2017; McCallum et al. 2017; Grimonprez et al. 2015; Li et al. 2004; Khodaparast et al. 2016). Thus far, no studies for chronic mouse vagus nerve interfacing have been reported.

The mouse cervical vagus nerve is challenging to interface with due to the small diameter of nerve (∼100 µm). Additionally, the electrode must 1) be flexible to account for animal movement and 2) have a small footprint to minimize nerve damage and ensure chronic survival. To satisfy these design criteria, microwire technology was explored. Our approach was to manually wrap the microwire around the exterior of the vagus nerve, like a cuff electrode. By placing the microwire and then insulating the electrode *in vivo* in place on the nerve, the flexibility and small footprint requirements were achieved. Additionally, microwires have a proven track record for successful chronic neural interfacing in both the CNS (Nicolelis et al. 2003; Worrell et al. 2008; Saxena et al. 2013; Karumbaiah et al. 2013; Prasad et al. 2012; Ward et al. 2009; Falcone et al. 2018; Schwarz et al. 2014) and the PNS (Gore et al. 2015; Srinivasan et al. 2015).

Developing a neural interface model in mice is critical due to the availability of transgenic and disease models, thus increasing the impact for future studies untangling mechanisms of various diseases. A focus was also placed on awake, non-anesthetized recordings, since anesthesia is known to alter neural activity (Brown et al. 2010; Michelson & Kozai 2018; Uhrig et al. 2018) and adjusting the isoflurane concentration can significantly impact vagus nerve activity (Silverman et al. 2018). Therefore, to increase spontaneous neural activity and to better model the clinical environment, an awake recording paradigm was developed.

In this study, two major innovations were developed to help dissect neural circuitry in disease models for bioelectronic medicine: 1) a novel wrappable microwire electrode to successfully interface with the mouse cervical vagus nerve chronically, and 2) the development of an awake cervical vagus nerve recording model in chronically implanted mice.

## Results

### Wrappable Microwire Electrode

Three Teflon-coated platinum microwires were soldered to a connector, and the distal end of the wire was de-insulated for neural interfacing (**Fig. 1A**). *In vivo*, each de-insulated microwire was individually tied in an overhand knot around the nerve and reinsulated using Kwik-Sil (silicone elastomer) (**Fig. 1B and C**). Spontaneous neural activity was acquired for a minimum of 30 minutes (**Fig. 2A and B**), and for the chronic preparation, viable CAPs were identified over time **(Fig. 2C**).

**Fig. 1:**
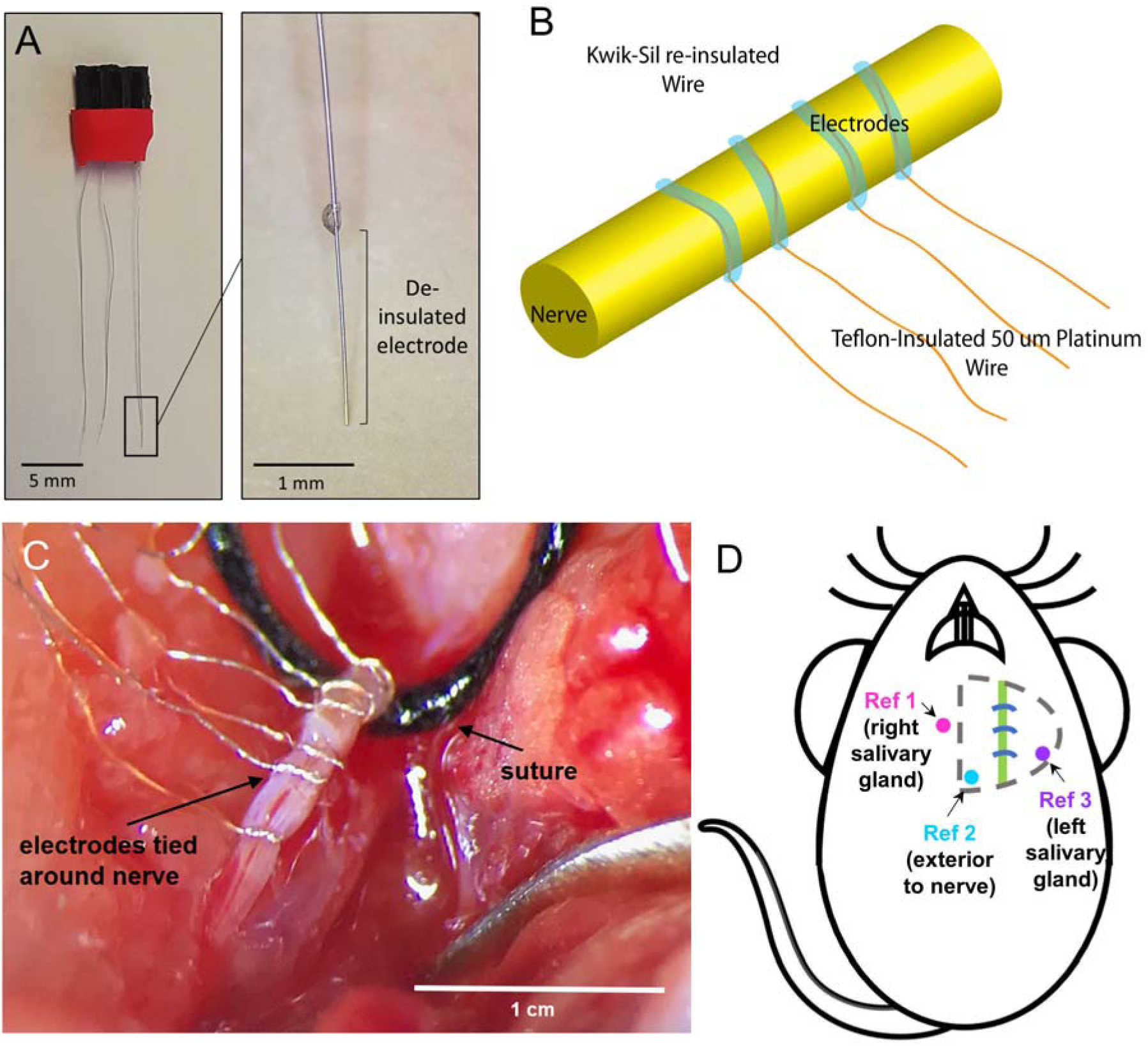
– Design concept and surgical implementation of the wrappable microwire electrode. **(A)** The microwires attached to the connector (left, scale bar = 5 mm) with a magnified view of the deinsulated portion of the microwire (right, scale bar = 1 mm). **(B)** Illustrative demonstration of the technique for wrapping the microwires around the vagus nerve and applying Kwik-Sil (blue). **(C)** Surgical example of microwires wrapped and knotted around the rat cervical vagus nerve (scale bar = 1 cm). **(D)** Illustration showing the placement of the three reference electrodes in relation to the vagus nerve (green) and the recording electrodes (blue). Reference 1 (pink) is on top of the right salivary gland, reference 2 (light blue) is situated near the vagus nerve, and reference 3 (purple) is on top of the left salivary gland.

**Fig. 2:**
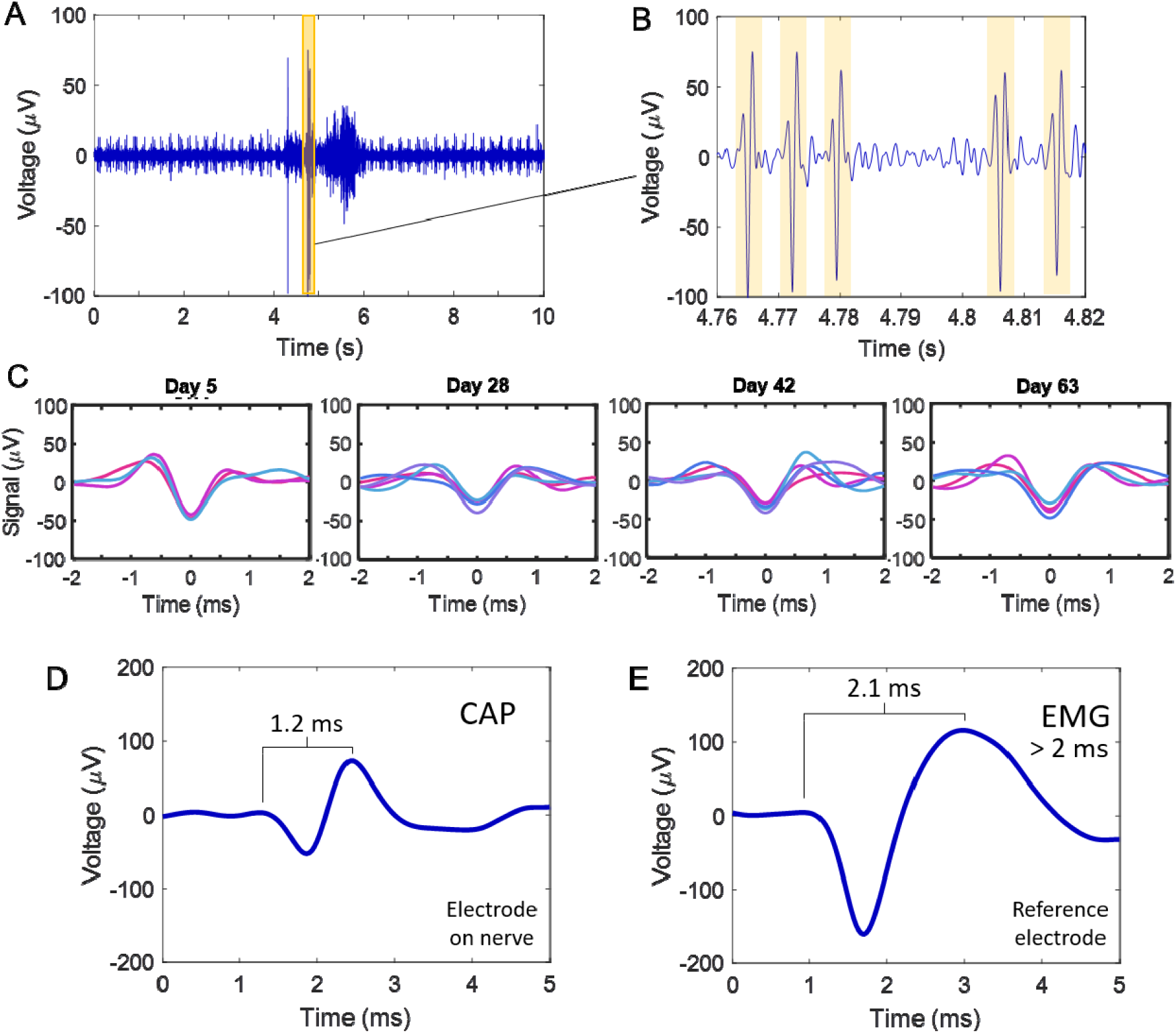
– Sample neural activity recorded from the cervical vagus nerve with the wrappable microwire. **(A)** A chronic bandpass filtered recording of the vagus nerve in an awake mouse. **(B)** Magnified view of the highlighted section from **(A)** with high SNR CAPs highlighted. **(C)** Representative mean waveforms (each color is a different sorted CAP) for all CAPs on a single channel from day 5 to day 63. Representative waveforms for **(D)** a CAP and **(E)** an electromyography (EMG). Note that the crest-to-crest wavelength of the EMG is longer than 2 ms and the peak-to-peak amplitude is larger than recorded CAPs.

### Acute Recordings

Three types of electrodes were compared in the acute model: 1) the novel wrappable microwire, 2) Cortec, and 3) Microprobes cuff electrodes. Electrodes were implanted on the left cervical vagus nerve in anesthetized mice and acute recordings were conducted. Spontaneous activity was recorded from the nerve for a minimum of 30 minutes. Recordings were analyzed with wave_clus to sort CAPs. Representative mean CAPs for each of the three electrodes are shown in **Fig. 3A**. Comparable mean peak-to-peak (P2P) amplitudes were found for isolated CAPs between the wrappable microwire electrode (n = 15 animals, 114 ± 11 μV), Microprobes (110 ± 28 μV) and Cortec (115 ± 7.9 μV) electrodes (**Fig. 3B**). For the SNR (**Fig. 3C**), no significant differences were found between the wrappable microwire electrode (7.58 ± 0.73) and the two commercial electrodes. The Cortec electrode (11 ± 2.8) had a significantly higher SNR than the Microprobes (5.78 ± 0.3) through percentile bootstrapping (*P* < 0.05).

**Fig. 3:**
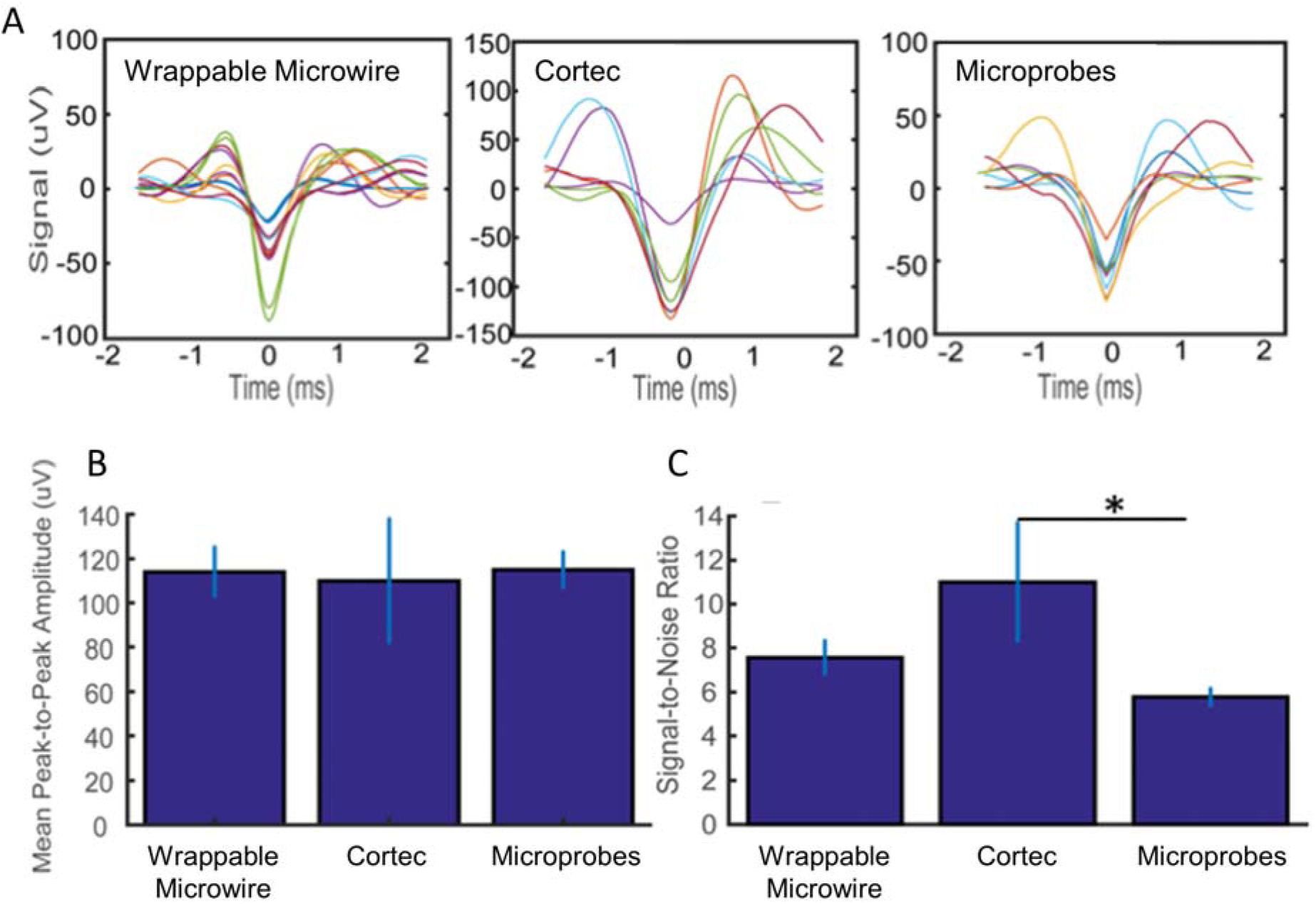
– Comparison of electrode types in an acute, anesthetized cervical vagus nerve recording model. **(A)** Representative mean waveforms for all CAPs on a single channel for a wrappable microwire, Cortec, and Microprobes electrode. Comparison of average **(B)** P2P voltages and **(C)** SNR showed that the wrappable microwire electrode was comparable in recording quality to Cortec and Microprobes commercial cuff electrodes (percentile bootstrapping: P < 0.05).

### Chronic Awake Recordings

An advantage of the wrappable microwire electrode is the ability to be implanted chronically. Spontaneous neural activity was recorded on awake mice (n = 8) for a minimum of 30 days, with a smaller cohort (n = 3) surpassing 60 days. Representative waveforms for mean CAPs on a single channel **(Fig. 2C**) demonstrate that viable CAPs were obtained over time. Three reference microwires had been implanted under both the left and right salivary glands and next to the nerve (**Fig. 1D**). Prior to the first recording session, each reference electrode was evaluated, and the reference with the lowest noise floor and lack of any EMG contamination (**Fig. 2D and E**) was selected.

The signal-to-noise ratio (SNR) was calculated by taking the P2P amplitude of CAPs and dividing by the P2P amplitude of the noise. The average SNR over time was plotted for each mouse (**Fig. 4A**, for individual SNR plots with +/- S.E.M., please see **Supplementary Fig. 2**) and remained above 1.3 (Ludwig et al. 2009). A two-way ANOVA of SNR revealed no significant difference over time (F(2,25) = 12.64, *P* > 0.05), and no significant difference between animals (F(2,4) = 12.85, *P* > 0.05). Additionally, *in vivo* electrode impedances were measured at 1 kHz during each recording session and normalized to the first day of measurement, typically 5 days after implantation (**Fig. 4B**). A two-way ANOVA of normalized impedance was not significant over time (F = 1.7, *P* > 0.05), but showed significant differences between animals (F = 3.69, *P* < 0.05). However, 5 out of 8 of the animals exhibited steady impedances from 30 to 60 days. There was a lack of correlation between impedance and SNR over time among all animals (**Fig. 4C**). For this data set, an R-squared value of 0.192 was calculated with a 1-degree polynomial test using the least absolute residual method. These results suggest that the wrappable microwire electrode is a reliable and repeatable electrode for vagus nerve interfacing over the time points examined.

**Fig. 4:**
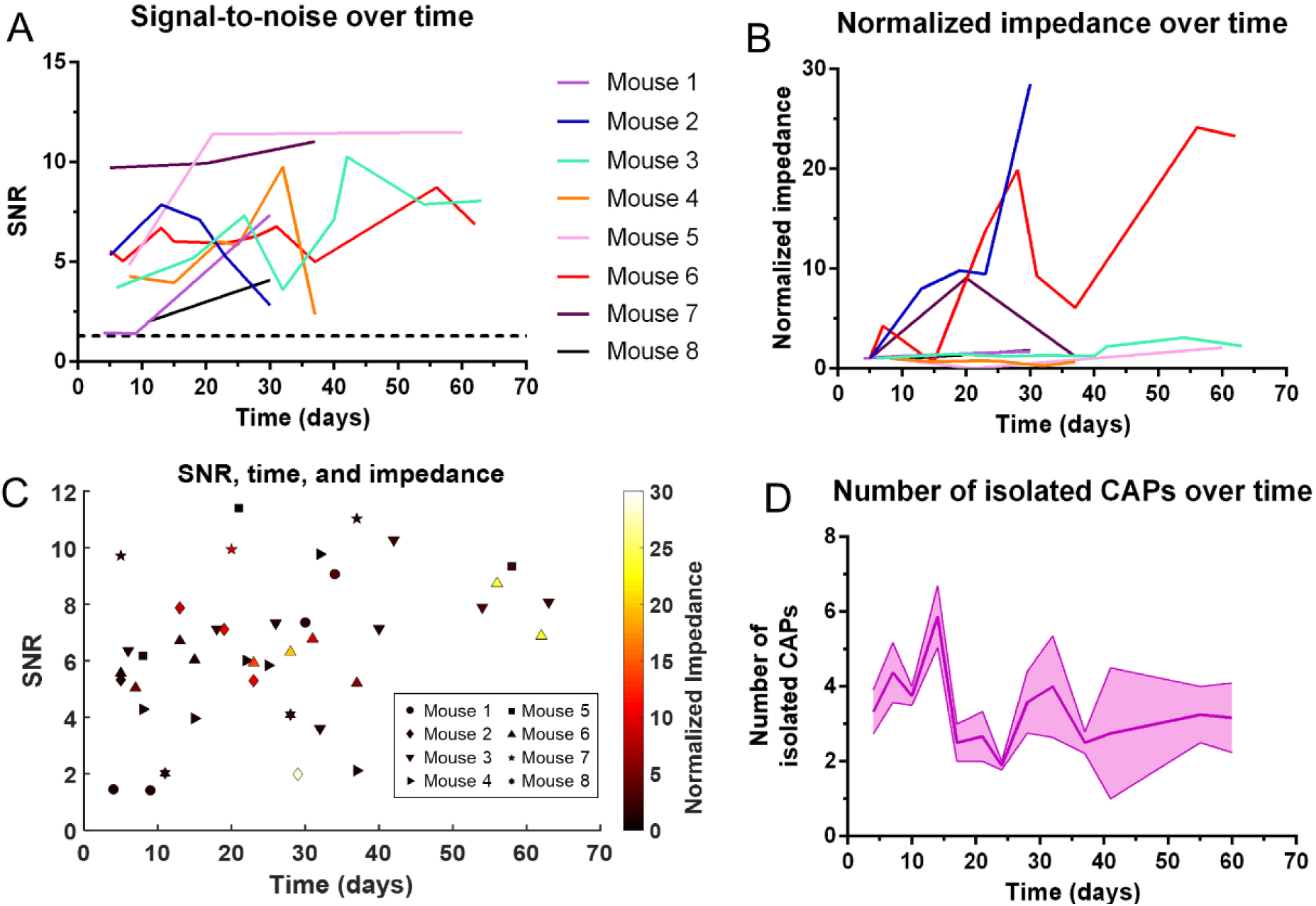
– Recording performance metrics of the chronically implanted wrappable microwire electrode. **(A)** Calculated average SNRs for each individual mouse were plotted over time. Acceptable SNRs (> 1.3, black dotted line) were obtained over all days for all animals, and no significant difference was observed between animals or across time (P > 0.05). **(B)** In-vivo impedance values at 1 kHz, normalized to the first recording session, for each individual mouse were plotted over time. There was no significant change over time for normalized impedance (P < 0.05). There was a significant difference between animals, however, it is noted that 5 out of 8 mice showed relatively stable impedances over time, indicative of a stable electrode-tissue interface. **(C)** Correlation plot of SNR (y-axis), time (x-axis) and normalized impedance (heatmap) for each individual mouse. No correlation of impedance and SNR were found (R-squared = 0.192). **(D)** Mean isolated CAPs plotted over time (+/- SEM) across all animals. No significant differences were found over time or across animals (P > 0.05) for the number of isolated CAPS.

The benefit of recording from multiple channels in a monopolar configuration is the ability to record signals with phase delay across channels. These phase delays can be used to calculate the conduction velocity of signals and perhaps distinguish between afferent and efferent pathways, if the same CAP is being captured on the channels (Caravaca et al. 2017). Examples of phase delays can be found in recordings obtained in the chronic awake model in the preliminary analysis. In **Fig. 5A**, the conduction velocity in box 1, calculated between channel 3 (gold) and either channel 1 (blue) or channel 2 (red), is 5 m/s; in box 2, the conduction velocity is 3 m/s, suggesting that the recorded CAP is from a A-δ fiber type.

**Fig. 5:**
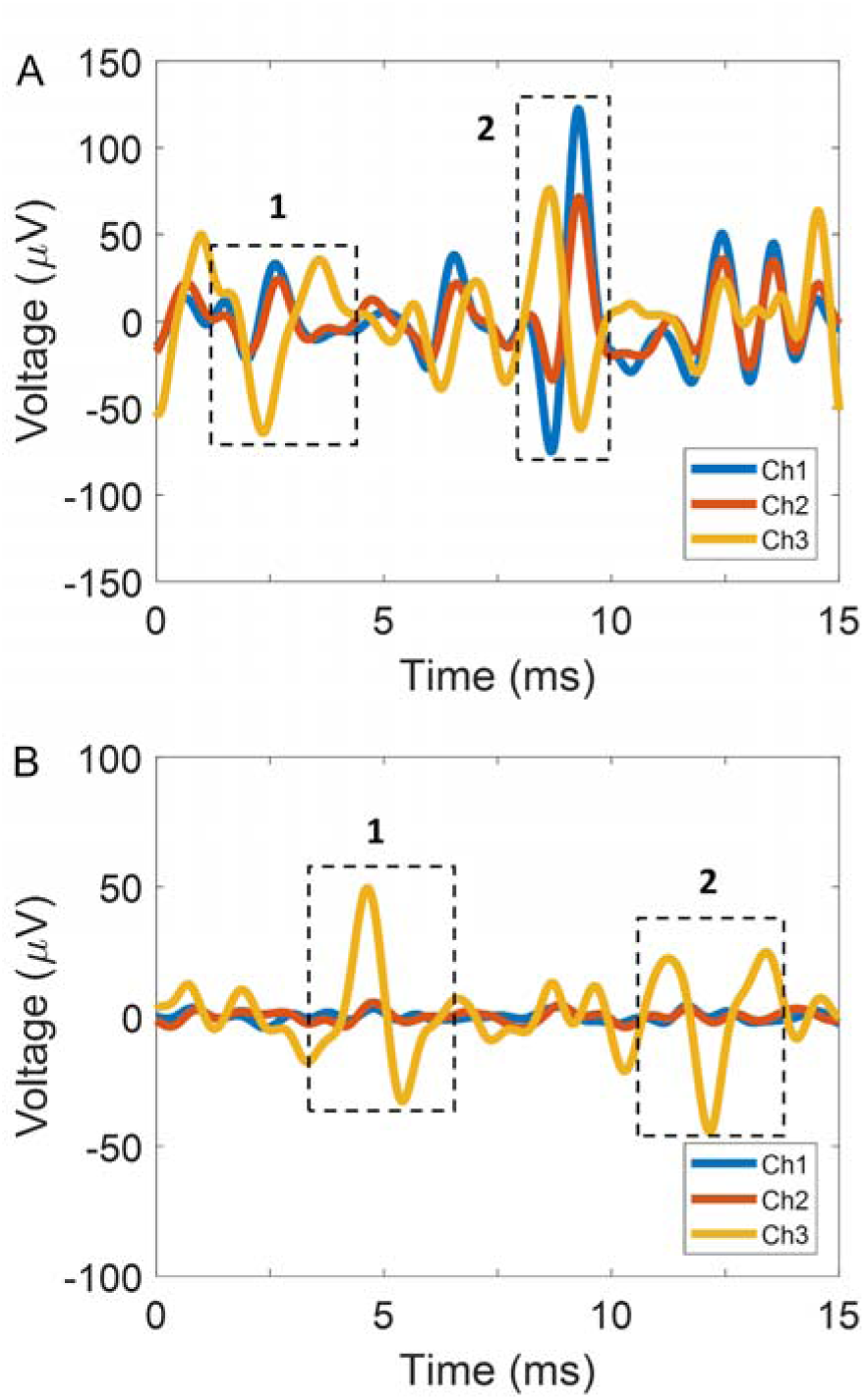
– The utility of having multiple electrode recording sites on the nerve was demonstrated. **(A)** Examples of recorded phase shifts between channel 3 (gold) and channels 1 and 2 (blue and red, respectively). The conduction velocity in box 1 is 5 m/s and in box 2, is 3 m/s suggesting the recording is from A-δ fibers. **(B)** Examples of CAPs being found on one channel (3, gold) but not on other channels, highlighting specific firing from isolated parts of the cervical vagus nerve in mice. These channels had similar impedance values.

In certain cases, spontaneous activity can be exclusively observed on one channel and not the others. In **Fig. 5B**, channel 3 (gold) displays CAP activity (as highlighted in box 1 and box 2), while minimal activity is observed on channel 1 and 2. This highlights the isolated firing patterns from specific parts of the nerve that exist in awake spontaneous activity, which is not apparent in evoked stimulation potentials from peripheral nerves (Gillis et al. 2018; Lee et al. 2017), where similar activity can be found across all the electrodes. Taken together, having multiple recording channels allows for the isolation of more specific nerve activity and allows for the elucidation of possible fiber types involved in CAP generation. This can be an invaluable tool when trying to understand mechanisms for bioelectronic medicine and contributions of specific fiber types in rodent disease models.

### Chronic mouse vagus nerve histology

After 30 days of implantation, the nerves from both the left (electrode implanted) and right (naive) vagus nerve were collected from a mouse (n = 1), selected at random. The tissue was paraffin embedded and transversely sectioned. H&E staining was conducted, and visible axons were counted within each nerve section (**Fig. 6C**). A student’s t-test showed no significant difference in between the number of axons for the naive (**Fig. 6B**) and the electrode implanted nerve (**Fig. 6A**) (average of 0.009 axons/µm^2^, t = 0.54, p > 0.05, n = 5 sections). It should be noted that axon diameters range in size from 1 µm to 5 µm.

**Fig. 6:**
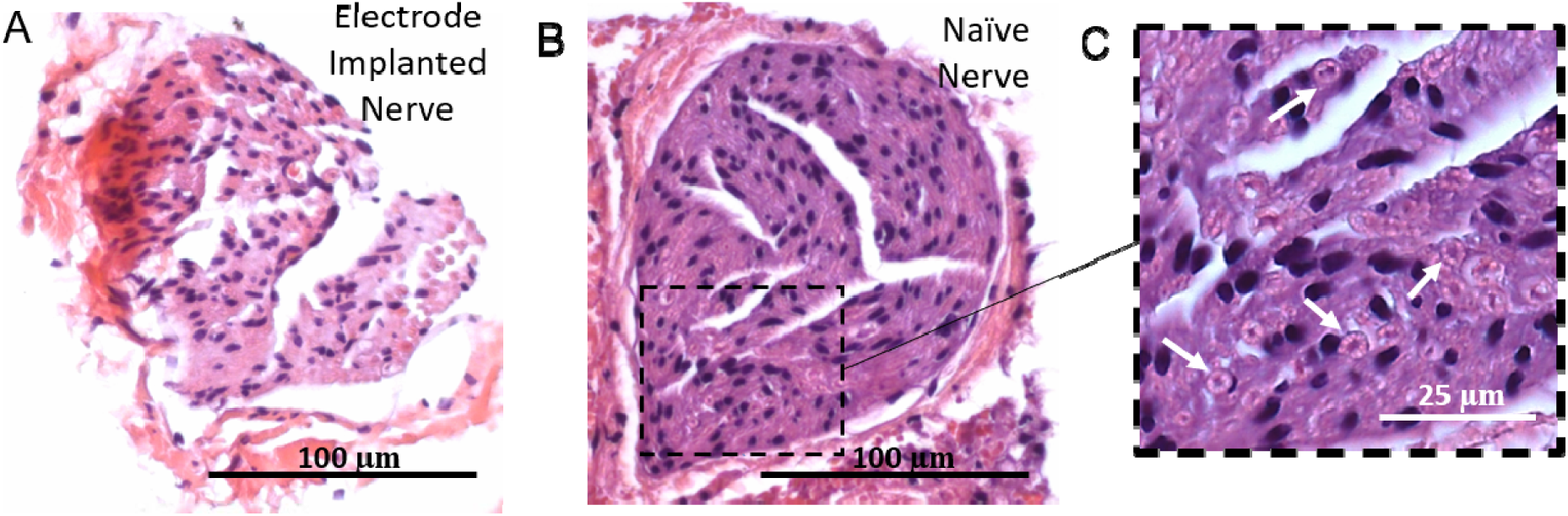
– Representative cross-sectioned H&E images of **(A)** an electrode implanted nerve and **(B)** a naive nerve (scale bar = 100 µm) post-30 days implantation. **(C)** Magnified portion of the naive nerve revealing visible axons (white arrows, scale bar = 25 µm).

## Discussion

To the best of our knowledge, this study is the first to validate a chronic, awake vagus nerve recording in mice. Vagus nerve recordings have been conducted under anesthesia for both acute (WOODBURY & WOODBURY 1991; Caravaca et al. 2017; Dias Da Silva et al. 2002; Krolczyk et al. 2008) and chronic models (Caravaca et al. 2017; Sauter et al. 1983). Others have chronically recorded from the peripheral sciatic nerve in awake rat models (Michoud et al. 2017; Srinivasan et al. 2015; Gore et al. 2015; Srinivasan et al. 2016; Musick et al. 2015; Vasudevan et al. 2017) and from the vagus nerve in anesthetized rat models (Khodaparast et al. 2014; Khodaparast et al. 2016; Somann et al. 2018), but these devices have not been translated to the mouse vagus nerve model. Here we demonstrated that this microwire based approach was effective in obtaining chronic neural recordings and identifiable CAPs from the cervical vagus nerve in mice.

To characterize chronic recording quality, SNR and impedances were measured *in vivo*. It has been postulated that changes in impedance *in vivo* directly correlate to signal changes due to the development of the glial scar (Polikov et al. 2005; Ward et al. 2009; McConnell et al. 2009). From this study, impedance fluctuations had no significant correlation with SNR. This trend has been confirmed in studies using intracortical electrodes from cortex recordings in the chronic setting (Barrese et al. 2013; Perge et al. 2013; Sohal et al. 2014; Suner et al. 2005; Kuo et al. 2013; Barrese et al. 2016).

Three *in vivo* checks can be performed to confirm that the electrode remains on the nerve: evoking bradycardia through stimulation on a weekly basis (if the electrode impedances are low enough for the compliance voltage of the stimulator), 2) ensuring that channels with similar impedances are detecting similar activity (in terms of SNR and waveforms), and 3) evaluating the spontaneous recording real time to confirm signals are CAPs and not EMG activity. These checks can be further confirmed with post-mortem nerve removal for histological examination.

Bipolar and monopolar recording paradigms were also compared, but no significant difference was found in terms of CAP quality. Bootstrapping between the number of obtained CAPs or SNR between a monopolar and bipolar amplifier setup showed no significant difference (percentile bootstrapping: *P* > 0.05). Monopolar recordings were therefore favored, as a bipolar derivation can always be implemented offline (with no data lost) through software subtraction of channels (e.g. through Matlab). Additionally, with monopolar recordings, it is possible to record phase shifts in CAPs and thus use conduction velocity to determine fiber types. This could be used to dissect pathways in bioelectronic medicine, for example, elucidating responses for the multiple fiber types that are involved in cytokine responses in the inflammatory reflex (Steinberg et al. 2016; Zanos et al. 2018; Silverman et al. 2018).

Although we report data from 8 mice that passed the 30-day mark, more animals were implanted (n = 11). In the initial cohort of implanted mice, headcap failure (n = 3) was failure mode for the devices, and the cause was the forces exerted on the connected headcap when the animal was restrained. However, the addition of skull screws (at least two are recommended) to anchor the dental acrylic and connectors led to successful recordings for over 60 days. Other animals at 30 days were sacrificed to check the electrode integrity and to ensure the electrode was still on the nerve for the animals.

An initial histological examination of the electrode implanted nerve showed that there were a similar number of fibers remaining in the nerve after the implantation period compared to the naive control. However, the resolution of the staining was restricted as a simple H & E stain was used to quantify the axon numbers in the nerve. Additionally, due to the small size of the nerve sections were often damaged during processing the tissue, leading to fractures in paraffin embedded slices. However, this discrepancy was similar for all nerve sections. Future studies will include a more detailed examination using immunohistochemistry and fluorescence to quantify the overall nerve structural integrity and the foreign body response (Srinivasan et al. 2015; Caravaca et al. 2017; Gillis et al. 2018; Christensen et al. 2016). Also, due to the small size of the nerve (∼ 100 µm) and indwelling axons (1 to 5 µm), methods to expand the fine tissue structures, such as expansion microscopy (Chen et al. 2015), might be beneficial.

Another benefit from the awake chronic preparation is the ability to compare spontaneous nerve activity between the anesthetized and awake states. We regularly observed increased spontaneous CAP activity with awake recovered animals (**Fig. 7C**). During anesthetized preparations, the respiratory modulation from the nerve (Steinberg et al. 2016; Silverman et al. 2018; Caravaca et al. 2017; Sevcencu et al. 2018) was reported as the dominant signal of the recording. In this study, the amplitude of the respiratory modulation visually decreased in awake real-time recordings compared to the anesthetized recordings (**Fig. 7A-C**). While recording in an awake animal removed the effects of anesthesia, motion artifacts and EMG contamination were introduced. Dual thresholding was implemented to capture CAPs and discard higher amplitude motion artifacts (approximately > 150 µV). Additionally, manual oversight was implemented to identify and remove myographic signals that were included in the threshold range. EMGs have a larger crest-to-crest period of > 2 ms compared to CAPs, as well as larger amplitudes (Lee et al. 2017; Xiang, Sheshadri, et al. 2016; Xiang, Yen, et al. 2016).

**Fig. 7:**
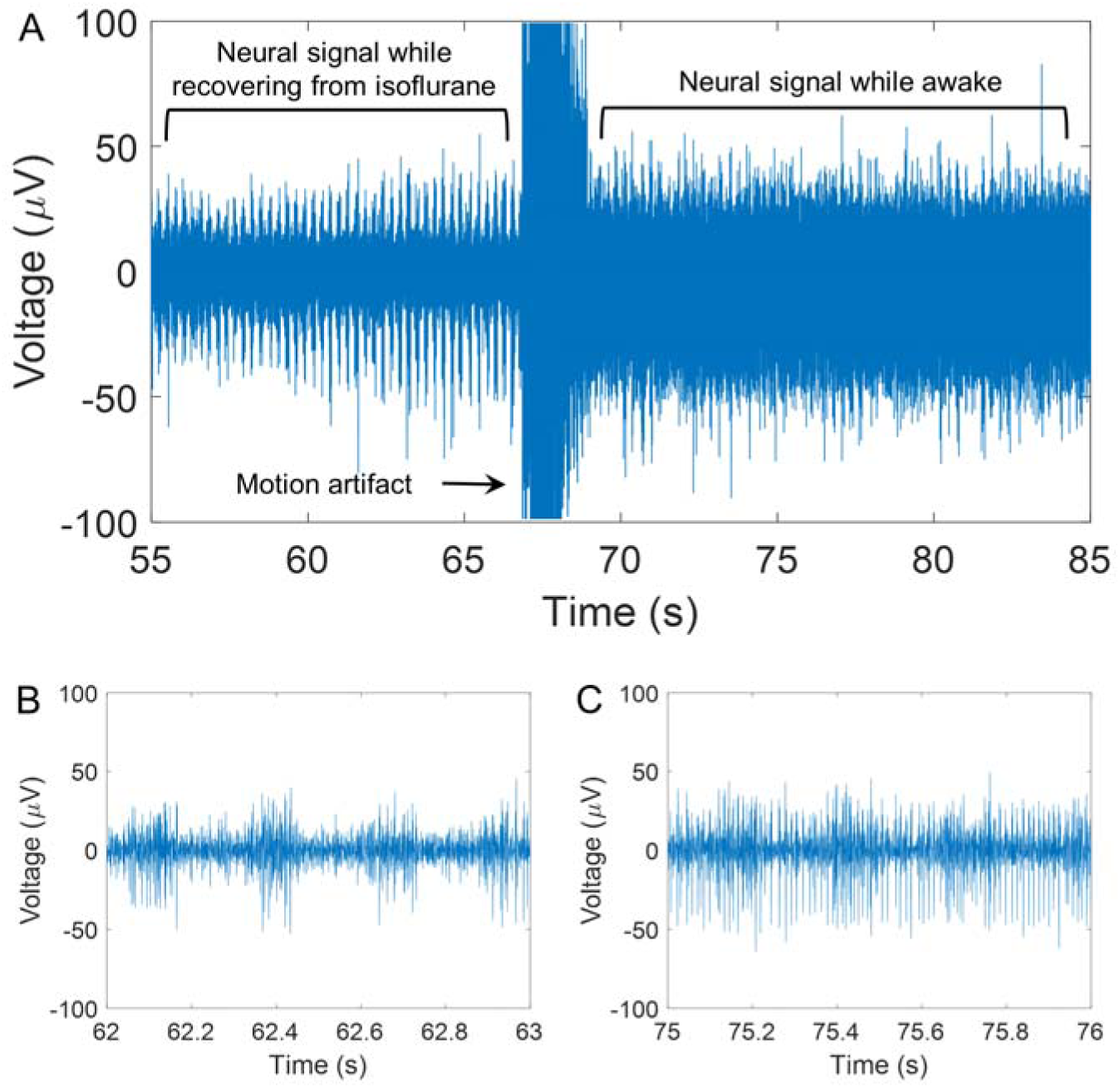
– Anesthesia effect on respiration signal. **(A)** Example recording of animal waking up after isoflurane induction. **(B)** Magnified section of the signal while the animal is recovering from isoflurane (prior to waking up). Note the repetitive respiration bursts. **(C)** Magnified section of the signal after the animal has recovered and is awake. Note the increase in spontaneous activity with respiration no longer the dominating signal.

For this study, awake recordings were conducted while the animal was physically restrained. It was noted that if the animal was stressed during the recording, the amplitude of respiratory modulation increased and was visually identified and associated with increased depth of breathing. Previous work has shown increasing the depth of lung inflation correlates with stronger vagus nerve activity (Zhang et al. 2008). Perhaps the increase in amplitude modulation in the anesthetized preparation could be explained with the increased depth of breathing elicited by inhalation anesthetics (e.g. isoflurane), although this does merit further investigation. Further, bursts of neural activity were observed to be associated with the animal defecating during recording. Characterization of basic physiological and autonomic functions in relation to the vagus nerve activity is necessary, particularly as the field moves towards recording in free-moving, awake, behaving models.

Microwires have been used to successfully interface with the CNS (Nicolelis et al. 2003; Worrell et al. 2008; Saxena et al. 2013; Karumbaiah et al. 2013; Prasad et al. 2012; Ward et al. 2009; Falcone et al. 2018; Schwarz et al. 2014) and the PNS (Gore et al. 2015; Srinivasan et al. 2015) in several different animal models. To further demonstrate the versatility of this electrode, the wrappable microwire was successfully implanted in the cervical vagus of a rat (**Supplementary Fig. 3**) for 105 days. As an example of recording quality, spontaneous activity on the rat cervical vagus nerve was recorded in an awake model (**Supplementary Fig. 3A**). Sortable CAPs were identified over the 105-day period (**Supplementary Fig. 3B, C**). While impedance increased over time (**Supplementary Fig. 3F**), viable SNRs were still recorded over the 105 days (**Supplementary Fig. 3D**), which were calculated from P2P amplitudes (**Supplementary Fig. 3E**). Similar to the results found in the awake mouse model, no correlation was found between impedance and SNR (R-squared = 0.038) or impedance and P2P (R-squared = 0.007). This initial feasibility result shows that it is possible to also use this method for rat peripheral nerves.

Indeed, this vagus nerve interfacing approach can be combined with neural technology for the brain (Chang et al. 2013). Head capping methods to anchor the external connector are the same for both intracortical and peripheral implants. In combination, this microwire electrode could be a useful tool to understand the contributions of the vagus nerve to brain function in models of disease (e.g. sepsis (Honig et al. 2016)), behavior (e.g. learning and memory (Yu et al. 2017; Stephenson-Jones et al. 2016)), and rehabilitation (Khodaparast et al. 2014).

The wrappable microwire electrode is a low-cost solution that can be adopted by many labs in the peripheral nerve interfacing community. Typical costs of commercial electrodes can range from $300 to $700 per device. At the time of the study, the cost of a 10-foot roll of microwire was $177 and connectors were purchased in bulk for $26 for 100 pins, equating to 33 electrodes. Typically, 2 inches of wire were used for each contact, which equates to less than $10 to fully assemble a three-contact electrode array. Another benefit of the wrappable microwire electrode is time for fabrication. Approximately 50+ electrodes can be fabricated in 1 week with a single person making electrodes. Even with salaries considered, the wrappable microwire electrode dramatically decreases both cost and lead-time compared to commercial counterparts.

Increasing the number of electrodes at the neural interface has the potential to improve specificity of neural recordings (Viventi et al. 2011; Rodger et al. 2008; Falcone & Bhatti 2011). This wrappable microwire electrode has the advantage of being a rapid prototype electrode at a low cost; however, the number of recording sites that can be placed around the nerve is constrained by the length of the excised nerve (typically 5 mm in the mouse). Therefore, this group is developing flexible, high density electrodes to chronically interface with the vagus nerve in the awake state (Caravaca et al. 2017; Li et al. 2017). These implants will increase the amount of information obtained from the nerve and also can be used to create unique stimulation patterns to potentially target different nerve fibers (Cassar et al. 2017).

Developing neural interfaces from flexible materials can reduce the foreign body response and allow the electrode to conform in an active implantation environment (Polikov et al. 2005). Many groups have been developing more mechanically compliant microwires using various polymeric materials (Kolarcik et al. 2015; Koppes et al. 2016; Park et al. 2017). This approach could be relevant to microwire interfacing groups to extend recording and potentially stimulation longevity. In addition, some of these polymeric microwire technologies have multimodal capabilities (e.g. light delivery for optogenetics (Park et al. 2017)), which could be useful for further dissection of circuits relevant to disease models in bioelectronic medicine.

Here we present a wrappable microwire electrode for chronic, awake recordings in the mouse cervical vagus model. Viable recordings were obtained from 30 to 60 days, which is a common timeframe for murine experiments. With this device, the high throughput demands of the preclinical environment can be met, better facilitating clinical translation for bioelectronic medicine for more informed disease progression and stimulation.

## Methods

### Electrode Assembly

Teflon coated platinum microwires (A-M systems, WA) were de-insulated at the proximal end and soldered to a 1 mm pitch header connector (Digikey, MN) (**Fig. 1A**). The soldered end was re-insulated with the use of two-part epoxy (Loctite, EA9017, USA) and once cured, the epoxy was covered with heat shrink (3M, USA). The distal recording end of the microwire was de-insulated prior to surgery and was often 1-2 mm in length.

### Electrode Implantation Surgery

This study and all experimental protocols were approved by the Institutional Animal Care and Use Committee (IACUC) at the Feinstein Institute for Medical Research, Northwell Health System (Manhasset, NY, USA), which follows the National Institute of Health (NIH) guidelines for the ethical treatment of animals. Male BALB/c mice (Charles River Laboratories 20-25 g) and male Sprague Dawley rats (Taconic Biosciences, 250-400 g) were used for interfacing with the cervical vagus nerve in acute and chronic models. Rodents were housed in a laboratory environment on a 12-h light and dark cycle at 25°C, with ad libitum access to food and water. Animals were anesthetized in an induction chamber on 3% isoflurane and then transferred to a nose cone at 2% isoflurane. Temperature was monitored rectally, and a heat mat was placed under the animal for the duration of the procedure. Puralube (Dechra, UK) was placed on the eyes, and hair was removed with Nair (Church & Dwight Co, NJ) and the surgical sites were sterilized with alternating povidone-iodine and alcohol pads prior to incision.

### Acute model

For acute studies, mice were fasted for 6 hours prior to surgery to limit recorded gut activity on the vagus nerve (Silverman et al. 2018). Following the induction and surgical preparation, the fasted rodent was placed in the supine position. A ventral midline cervical incision was then performed, and subcutaneous tissues were retracted laterally. The salivary glands were bluntly separated and retracted away from the operative field. The left neurovascular bundle was revealed, and the cervical vagus nerve was then dissected away from the vasculature and isolated with a suture.

Three types of electrodes were compared acutely: 1) a micro-cuff sling with two platinum-iridium electrodes (500 x 100 µm recording site area) and silicone insulation (Cortec, Germany), a custom-made cuff electrode with platinum electrodes (5 electrodes, spaced 0.5 mm apart, 12 µm diameter wire) encased in a polyimide tube and silicone (Microprobes, MD), and 3) the wrappable platinum microwire electrode (3 electrodes, spaced ∼1 mm apart, 50 µm diameter wire) encased with Kwik-sil insulation applied during surgery. The first two were commercially available cuff electrodes. For the wrappable microwire electrode, each of the three de-insulated platinum microwires was individually tied in an overhand knot around the circumference of the nerve (**Fig. 1B and C**). A minimal application of Kwik-sil, an adhesive silicone elastomer (World Precision Instruments, FL), was placed around the wires for insulation (**Fig. 1B**) with the use of an insulin syringe (27G). This was to prevent signal loss due to shunting and to ensure that we only recorded activity from the nerve. After electrode implantation, a reference wire was then placed under the contralateral salivary gland as described previously (Silverman et al. 2018; Caravaca et al. 2017; Steinberg et al. 2016).

### Chronic model

Following induction and surgical preparation, the non-fasted rodent was placed in the prone position. A midline incision was performed, and the skin was retracted laterally to expose the skull. Tissue was removed from the skull, and the area was cleaned with 3% hydrogen peroxide, followed by 2 precision screws mounted on the skull. The screw location was marked with a sterile marker and self-tapping stainless-steel bone screws (1.17 mm diameter) were drilled into the skull. A thin layer of dental cement (A-M systems, WA) was added to secure the skull screws.

Next, the animal was switched to the supine position for a ventral midline cervical incision and retraction of subcutaneous tissues. The electrode wires were subcutaneously tunneled from the skull towards the midline cervical incision. As with the acute preparation, the salivary were separated and the cervical vagus nerve was dissected away from vasculature. Three (for mice) or six (for rats) de-insulated platinum microwires were tied around the circumference of the nerve with the remaining wire cut down to size using micro-scissors (**Fig. 1C**). Ideal electrodes spacing is a minimum of 1 mm to maintain mechanical independence for each recording site and Kwik-sil was then applied to provide insulation for the electrode. Additionally, Kwik-sil mechanically isolates each wire, minimizing possible tethering forces on the nerve, which is a phenomenon that occurs with the bulk cable from commercial cuff electrodes. The reference microwires were placed under both the left and right salivary glands and next to the nerve (**Fig. 1D**). The cervical area was sutured, and the animal was placed back in prone position. The connectors were fixed to head of the mouse with dental cement and the loose skin was tightened around the acrylic headcap with sutures and secured further with VetBond (3M, MN, USA). The animal was injected with buprenorphine (sub-q) and recovered on a heating pad.

### In Vivo Electrophysiology

Acute recordings were conducted under anesthesia immediately after placing the electrode around the nerve. For chronic studies, vagus nerve recordings were performed weekly (starting on day 7 after surgery) on awake animals in a restraining tube (Braintree Scientific, MA) to reduce motion artifact and give easy access to the tail vein. Electrophysiological data was acquired with the Intan Technologies RHD2000 Recording System (Intan Technologies, USA). Monopolar recordings were sampled at 30 kHz and a 160 to 3000 Hz bandpass filter was applied. Spontaneous activity was acquired for a minimum of 30 minutes (**Fig. 2A and B**). For the chronic studies, prior to the first recording session, each reference electrode was evaluated, and the “best” reference electrode was chosen for the recording. The lowest noise floor and lack of any EMG contamination (**Fig. 2E**) attributed to the choice of reference. The selection of the reference electrode was kept constant throughout the chronic characterization period. Impedance values at 1 kHz were measured at the interfacing electrodes relative to the reference for every recording session using the Intan Recording Controller system and bundled software.

### Offline Signal Analysis

Recordings in the form of .RHD files were imported into Matlab (2016a, Mathworks, USA) and converted for CAP analysis. Dual thresholds were used: the first for CAP detection and the second, a larger threshold, for artifact rejection (e.g. motion and often EMG). Batch sorting in wave_clus (Quiroga et al. 2004) was then conducted (**Fig. 2C, Supplementary Fig. 1**). Sorted CAPs were manually inspected to reject any non-neural activity, and additional criteria for sorted CAPs included: 1) acceptable inter-CAP-interval histograms, 2) a minimum CAP cluster over 100 waveforms, and 3) acceptable CAP firing rate (**Supplementary Fig. 1**).

Representative waveforms in **Fig. 2D and E** demonstrate the differences between CAPs and EMGs. As shown, the shapes of CAPs and EMGs can be similar; however, the positive crest-to-crest duration of a CAP should be under 2 ms when using a negative threshold detection (Xiang, Yen, et al. 2016; Xiang, Sheshadri, et al. 2016; Lee et al. 2017). Any sorted waveforms that were over 2 ms were discarded. Further, EMG amplitudes were typically larger than those of CAPs (> -200 µV P2P amplitude), and this was used as an additional metric to discard misidentified CAPs.

Isolated CAPs were assessed in terms of recording quality by calculating the peak-to-peak amplitude (P2P) and signal-to-noise ratio (SNR) for both the acute and chronic awake recordings (Ludwig et al. 2009). The mean P2P amplitude of the sorted waveforms was calculated. For the SNR, the mean P2P was divided by the average P2P amplitude of the noise floor, obtained from the noise floor of the 160 to 3000 Hz band pass filtered signal. Other common methods used for SNR calculation where the noise value is derived from the signal (Suner et al. 2005; Sohal et al. 2014; Vetter et al. 2004) are not appropriate for CAP analysis as the activity being recorded is more multi-unit activity rather than single units, since multiple fibers contribute to a CAP response. Therefore, any SNR measures looking at the tightness of all overlain waveforms relative to the mean waveform will have a low SNR on principle.

### Nerve Histology

After 30 days, a mouse was sacrificed with an overdose of isoflurane (5%). The electrode and left vagus nerve were removed, along with the right vagus nerve as a naive control and stored in formalin at 4°C to preserve the tissue for subsequent histology. The electrode was removed, and the tissue was then embedded in paraffin and sectioned transversely at 5 µm. Slides were incubated in xylene and serially dehydrated in ethanol. Haemotoxylin and Eosin (H&E) were applied separately to the slides, which, following incubation, were rinsed and mounted. Colored, bright field images were taken at 40x and 63x on an inverted microscope (Zeiss Apotome, Germany). Distinguishable axons were counted and compared between the naive nerve and electrode implanted nerve.

### Statistical Tests

All statistical tests were performed in the Matlab environment.

To compare the recorded responses between electrodes for the acute recordings, percentile-bootstrapping was used because of unequal sample sizes: the number of CAPs per recording session differed for each electrode type and animal (Sohal et al. 2014). CAP events from the full dataset were drawn randomly (with replacement) to create surrogate datasets with the same number of events as the real data and distributed evenly between all electrode types. This procedure was repeated 10000 times to estimate the expected difference in each measure under the null hypothesis of no difference between electrode types. The null hypothesis was rejected if the observed difference fell outside the 2.5–97.5% percentile range of the bootstrapped distribution.

The output metrics analyzed for electrophysiology were SNR and impedance. To evaluate change over time or inter-animal variability, a two-way analysis of variance (ANOVA) was conducted for each metric. For comparing the number of counted axons between implanted and normal nerve sections for the H&E histology, a student’s t-test was performed. For all tests, p-values less than 0.05 were deemed significant.

## Acknowledgments

We would like to thank members of the Center for Bioelectronic Medicine (Theodoros Zanos, Stavros Zanos, Yousef Al Abed) and Tracey lab (Eric Chang, Harold Silverman and Tea Tsaava) for insightful comments and discussions. We would also like to thank Pato Huerta for comments and suggestions regarding electrophysiology. This work was funded through internal Feinstein Institute funds.

## Author Contributions

HSS, LR, and CEB conceptualized this study. HSS and LG invented the electrode design for this study. LG, DP, TL, and HSS constructed the devices used in this study. LG, MS, TL, and HSS performed acute and chronic surgeries. JDF, TL, LG, and HSS collected neural data. JDF and HSS performed data analysis and interpretation. JDF and HSS drafted the original article. HSS, JDF, CEB, LR, and MS performed critical revision of the article.

